## Supplemental Figure 1-8 and Table 1 for "Acetylcholine modulates prefrontal outcome coding during threat learning under uncertainty": Tu_Supplemental_Final_Biorxiv.pdf

### SUPPLEMENTAL INFORMATION

Supplemental Fig S1-S8……………………….…………………………………………… 2
Supplemental Table S1

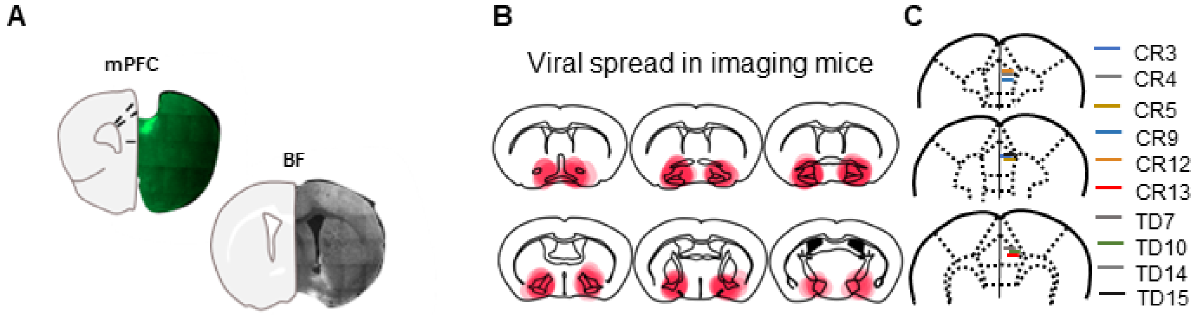


D

| Stage | Pre-training (day 1-5) | | | | | Probabilistic (day 6-10) | | | | | Random (day 11-15) | | | | |
|---|---|---|---|---|---|---|---|---|---|---|---|---|---|---|---|
| Session | PT1 | PT2 | PT3 | PT4 | PT5 | P1 | P2 | P3 | P4 | P5 | R1 | R2 | R3 | R4 | R5 |
| *P(Threat\|ht-p)* | 0 | 0 | 0 | 0 | 0 | 75 | 75 | 75 | 75 | 75 | 50 | 50 | 50 | 50 | 50 |
| P(Threat\|lt-p) | 0 | 0 | 0 | 0 | 0 | 25 | 25 | 25 | 25 | 25 | 50 | 50 | 50 | 50 | 50 |
| Orange LED | | | | | | | | | | | | | | | |

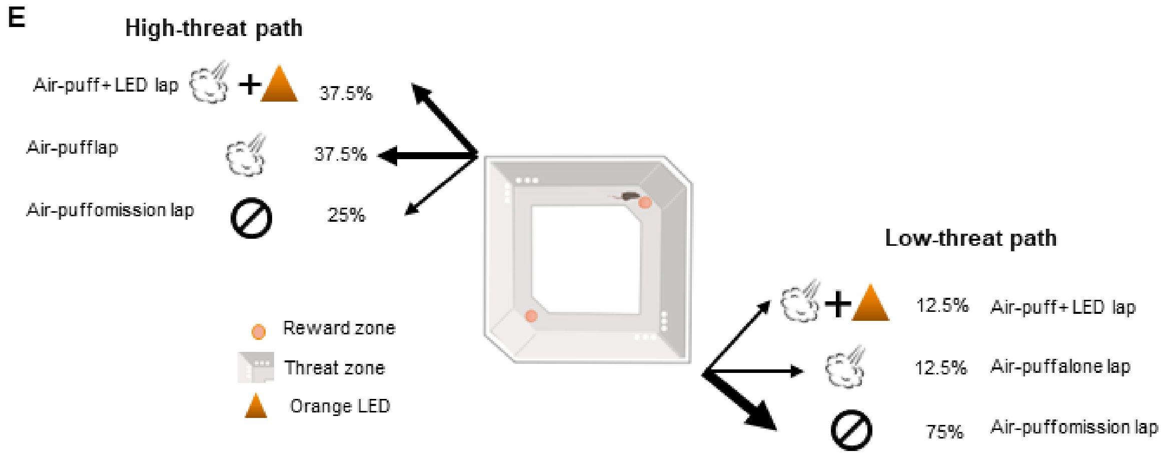


**Fig S1. Histological verifications and experimental design of the calcium imaging with optogenetic manipulation.** Related to Fig 1.
(**A**) Images showing viral spread (green) and GRIN lens location in the medial prefrontal cortex (mPFC) and viral spread (white) in the basal forebrain (BF). (**B**) Schematic representations of virus spread in the BF. Each red translucent area indicates the spread in each mouse infused with AAV5-Syn-Flex-ChrimR-tdTomato. (**C**) GRIN lens location in the prelimbic region of mice expressing ChrimsonR-tdTomato (CR) and tdTomato alone (TD). (**D**) Task schedule of the imaging experiment. The green color indicates the day on which calcium imaging was conducted. The orange color indicates the day on which LED stimulation was applied to the imaging window during a subset of air-puff delivery. (**E**) Schematic representation of three lap types. On both paths, ~50% of air-puff delivery was accompanied by LED stimulation.

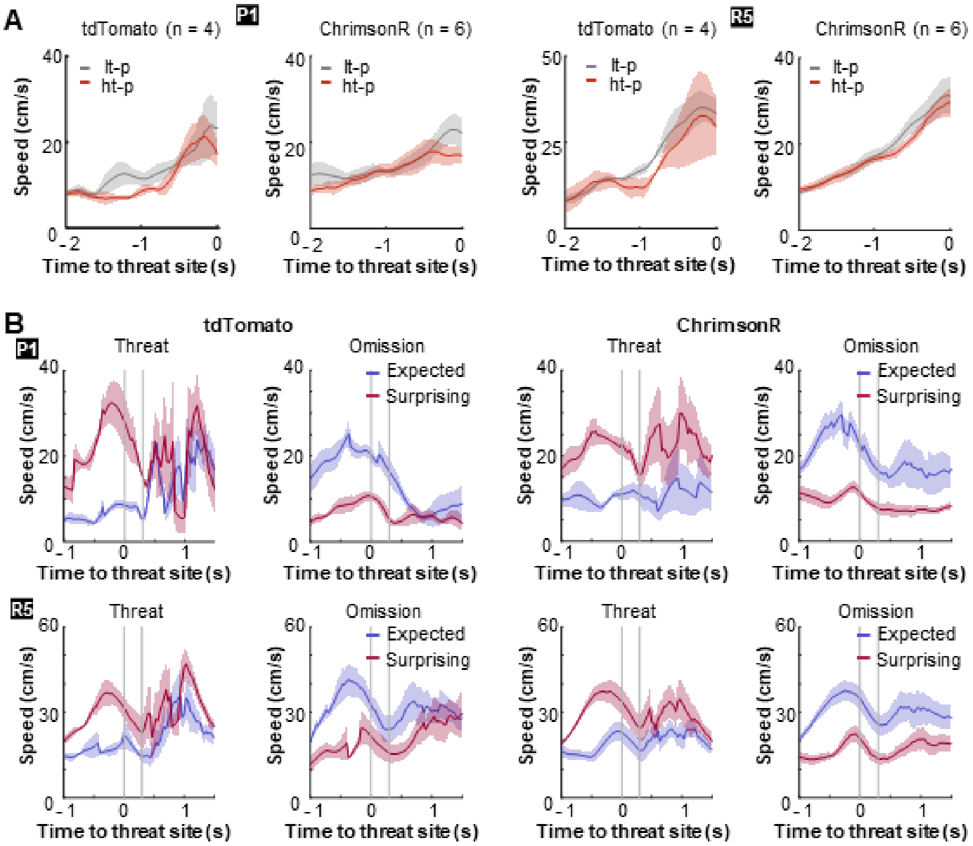


**Fig S2. Differentiation of movement speed depending on threat probabilities and expectations.** Related to Fig 1.

(**A**) The movement speed of mice while they approached the threat sites in the first session of the probabilistic stage (P1) and the last session of the random stage (R5). The running speed was averaged separately for the high-threat path with a 75% chance of air-puff delivery (ht-p) and the other low-threat path with a 25% chance of air-puff delivery (lt-p). Neither group differentiated the speed toward the threat site between the two paths in these sessions. Lines indicate the mean, and shaded areas indicate s.e.m. (**B**) Expectation-dependent differentiation of reactions to outcomes in P1 and R5. Laps were categorized into two types, fast and slow laps, based on the speed toward the threat sites (median split in each session). In each type, laps were further categorized into two types depending on whether mice received an air puff (threat) or not (omission). Fast laps with threats and slow laps with omissions were used to examine mice's reaction to surprising outcomes (red). Conversely, slow laps with threats and fast laps with omissions were used to examine their reaction to expected outcomes (blue). In R5, both groups moved faster after surprising threats than expected threats. In parallel, they took longer to accelerate after surprising omissions than expected omissions. Such differentiation was present only weakly in P1. Lines indicate the mean, and shaded areas indicate s.e.m.

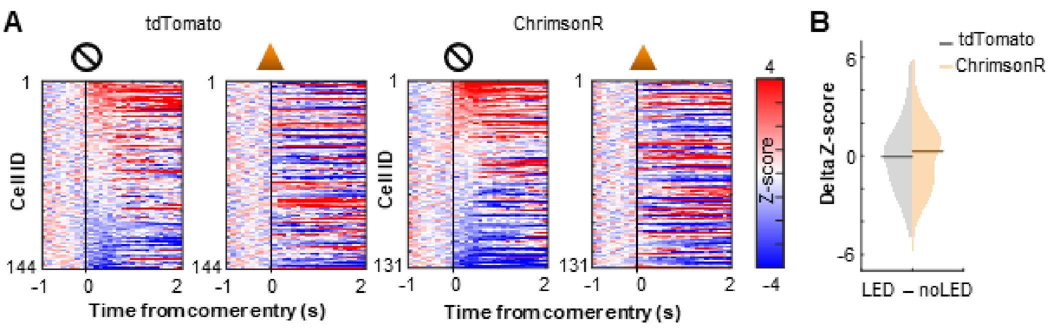


**Fig S3. Effects of cholinergic terminal stimulation on location-selective activity of PL cells.** Related to Fig 2.

(**A**) Pseudocolor plots showing PL cell activity during one pre-training session during which mice ran on the maze without air-puff delivery. The activity was z-normalized with the mean and standard deviation of the activity during a 1-second window before the entry to the corner of the maze that would become a threat site in subsequent days. In half of the laps, LED stimulation was applied to the imaging window upon the corner entry. The activity was normalized separately for laps with LED stimulation (right, LED) and the others without (left, noLED). Cells were sorted by the activity at the corner in noLED laps. (**B**) The distribution of the change in cell activity evoked by LED stimulation. Horizontal lines show the median. Z-normalized activity was averaged over a 500-msec window starting from the corner entry in each lap type. The average activity in noLED laps was subtracted from the average activity in LED laps. The degree of activity change with LED stimulation did not differ depending on the expression of ChrimpsonR in cholinergic terminals (Kolmogorov-Smirnov test, $p = 0.237$).

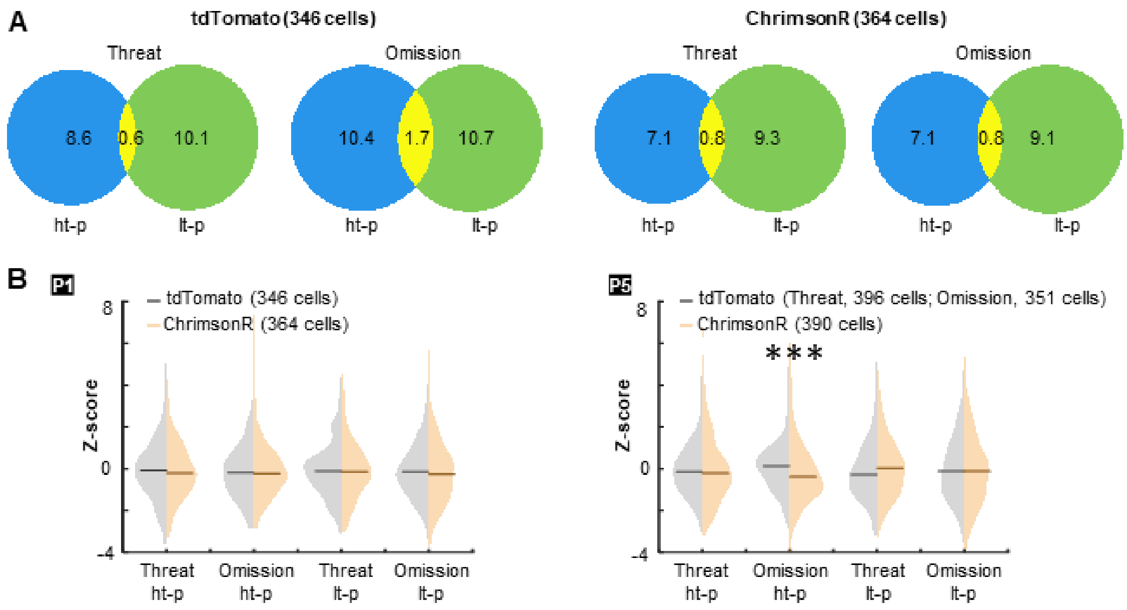


**Fig S4. Learning-dependent changes in outcome-related activity of PL cells.** Related to Fig 3.

(**A**) Venn diagrams depicting the proportion of cells responding to threats and their omission on the high-threat path (ht-p; blue) and the low-threat path (lt-p; green), as well as their overlap (yellow) during the first session of the probabilistic stage. In both groups, outcomes on two paths were represented by distinct groups of cells. The numbers indicate the proportions of cells in each category. (**B**) The distribution of the cell activity evoked by threats and their omission. Horizontal lines show the median. Z-normalized activity was averaged over a 500-msec window starting from the threat site entry in each lap type. During the first session (P1, left), two groups showed comparable magnitude of responses to threats and their omission on the ht-p and lt-p paths (Kolmogorov-Smirnov test, threat on ht-p, $p = 0.731$; omission on ht-p, $p = 0.682$; threat on lt-p, $p = 0.404$; omission on lt-p, $p = 0.142$). In the last session (P5, right), omission-evoked activity on the ht-p was weaker in ChrimsonR-expressing mice than no opsin control mice ($p < 0.001$). The group difference in other activity types did not reach statistical significance ($\alpha = 0.05/4$; threat on ht-p, $p = 0.054$; threat on lt-p, $p = 0.017$; omission on lt-p, $p = 0.409$). One no opsin mouse was removed from the analysis on omission-evoked activity in P5 because it did not experience omission on the ht-p.

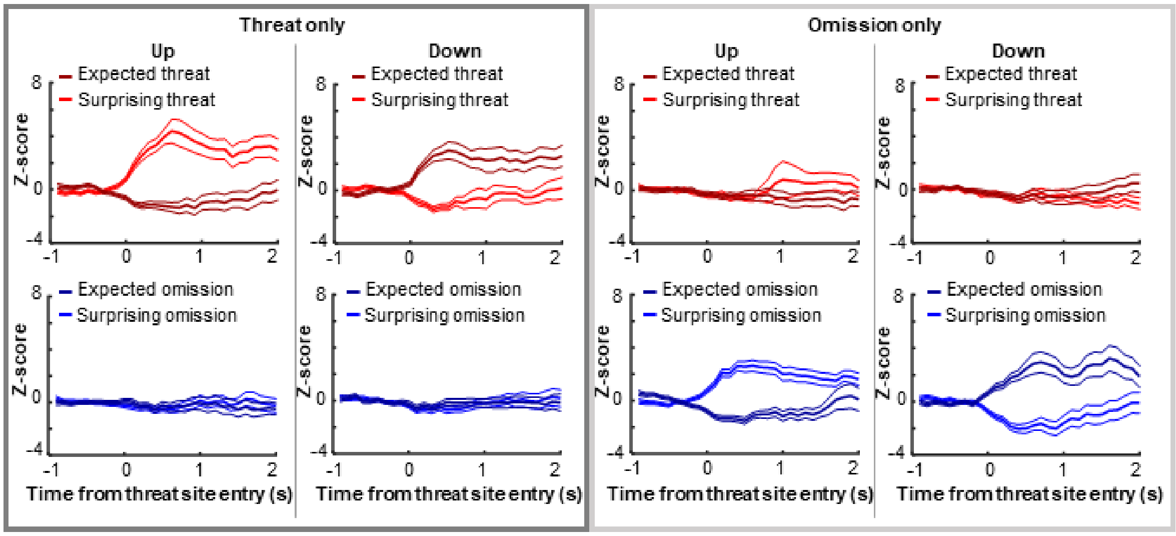


**Fig S5. Expectation-dependent differentiation of outcome-related activity of PL cells.** Related to Fig 4.

The average z-normalized activity across cells with different types of activity differentiation (mean ± s.e.m.). A cell was selected when its differential activity for threats or their omission was greater than 2 (Up) or smaller than -2 (Down). Some cells differentiated their responses to threats, but not their omission (Threat only). In parallel, other cells differentiated their responses to threat omission, but not theats (Omission only).

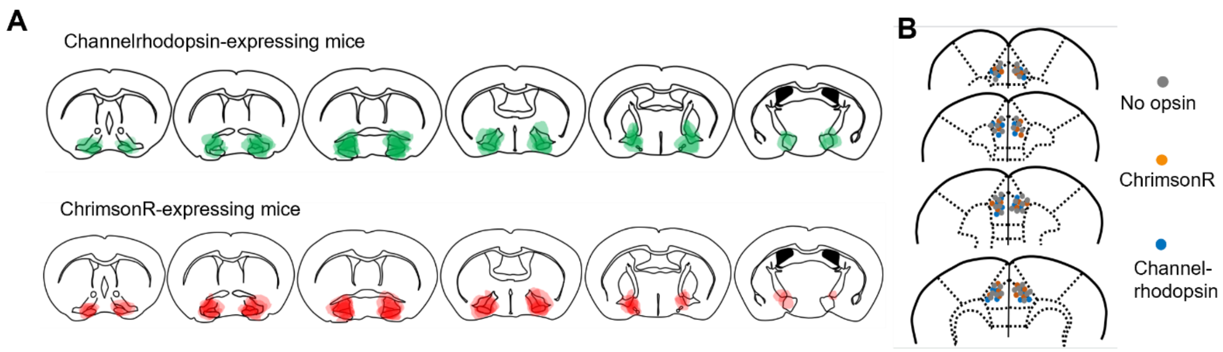


**Fig S6. Locations of viral spread and optic fiber implants in mice underwent bilateral optogenetic manipulations.** Related to Fig 6.

(**A**) Schematic representations of virus spread in the basal forebrain for mice expressing channelrhodopsin-EYFP (green, top) or ChrimsonR-tdTomato (red, bottom). Each translucent area indicates the viral spread in each mouse. (**B**) Schematic representations of optic fiber location (one dot/each fiber placement in each mouse).

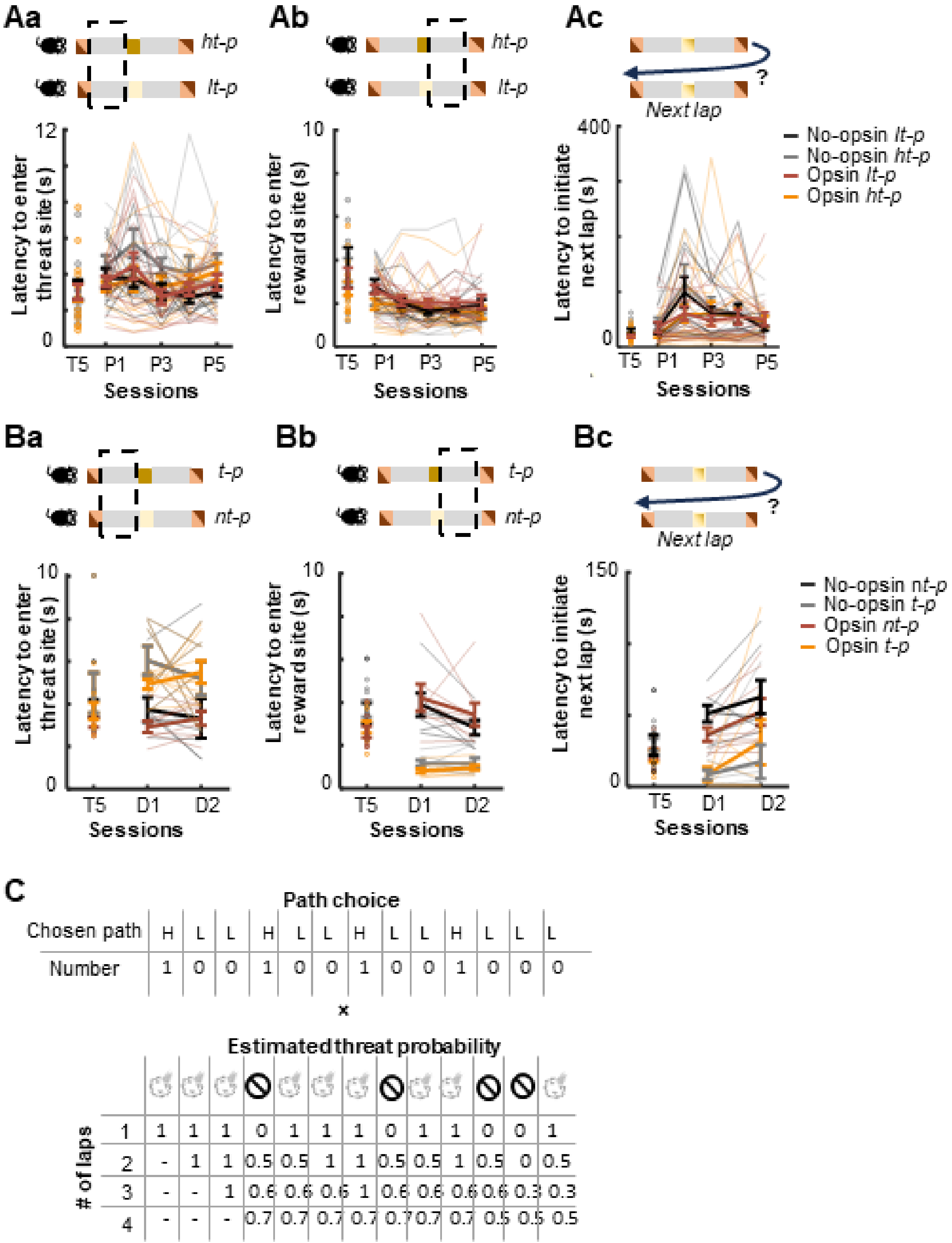


**Fig S7. Movement patterns and the design of correlation analysis.** Related to Fig 6.

(**A**) Movement patterns on the high- (ht-p) and low-threat path (lt-p) during the last pre-training session (T5) and all probabilistic sessions (P1-P5). No opsin, n = 11; Opsin, n = 15. a. The time taken from the reward site exit to the threat site entry. b. The time taken from the threat site exit to the reward site entry. c. Time in the reward site before initiating a subsequent lap. (**B**)

Movement patterns on the path with threats (t-p) and path without threats (nt-p) during the last pre-training session (T5) and all deterministic sessions (D1-2). a. The time taken from the reward site exit to the threat site entry. b. The time taken from the threat site exit to the reward site entry. c. Time in the reward site before initiating a subsequent lap. (**C**) chematic explanation of the correlation analysis in Figure 6F. The path that each mouse took in each lap was converted to 1 and 0 for the high- and low-threat path, respectively. In parallel, threat probabilities were estimated based on threat occurrence across different numbers of trials. We then calculated a Pearson correlation coefficient between the path choice and estimated threat probability.

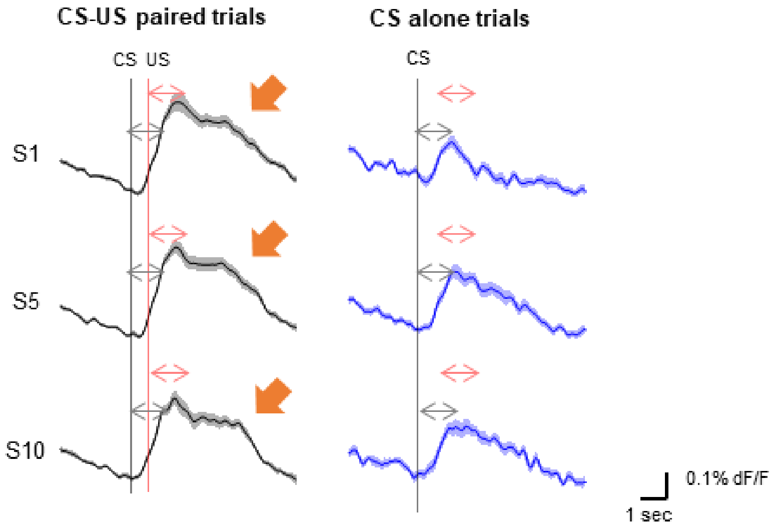


**Fig S8. Lack of omission-evoked activation of cholinergic terminals in the PL** Related to Fig 4 and discussion.

In our previous study[58], we conducted photometric recording from cholinergic terminals in the PL while mice associated an auditory conditioned stimulus (CS, 100 ms) with an aversive unconditioned stimulus (US, 100 ms) over a 500-msec stimulus interval. In each session, mice received 125 CS presentations. 80% of the CS was paired with the US (CS-US paired trials), while the remaining 20% of the CS was presented alone (CS-alone trials). These trials were inter-mixed and presented in a random order. Over the course of ten daily sessions (S1-S10), mice developed conditioned responses to the CS, indicating the formation of CS-US association[58]. From S1, cholinergic terminals were strongly activated by the CS (grey arrows) and the US (red arrows). Due to the slow kinetics of calcium indicator (GCaMP6s), the responses to the US overlapped with those to the CS, resulting in a longer peak latency and longer response duration (orange arrow) in CS-US paired trials than CS-alone trials. With learning, the magnitude of CS-evoked components became larger in both CS-US paired and CS-alone trials. However, the response latency or duration was never extended in CS-alone trials, indicating a lack of components evoked by surprising US omission. Thus, cholinergic terminals in the PL convey the presence of threat and threat-predictive cues, but not surprising threat omission.

**Table S1 Summary of the number of imaged cells in each mouse**

| Group | Mouse ID | P1 | P5 | R5 |
|---|---|---|---|---|
| tdTomato | M7 | 114 | 120 | 131 |
| tdTomato | M10 | 31 | 26 | 25 |
| tdTomato | M14 | 173 | 205 | 162 |
| tdTomato | M15 | 28 | 47 | 63 |
| ChrimsonR | M3 | 130 | 122 | 131 |
| ChrimsonR | M4 | 28 | 20 | 14 |
| ChrimsonR | M5 | 121 | 146 | 137 |
| ChrimsonR | M9 | 22 | 21 | 16 |
| ChrimsonR | M12 | 45 | 61 | 66 |
| ChrimsonR | M13 | 18 | 20 | 21 |
